# Contributions of T helper 9 cells in endometriosis-associated inflammation and lesion growth

**DOI:** 10.1101/2025.04.04.647212

**Authors:** Alison McCallion, Katherine B. Zutautas, Danielle J. Sisnett, Priyanka Yolmo, Harshavardhan Lingegowda, Asha K. Ravishanker, Dan Vo Hoang, Chandrakant Tayade

## Abstract

Endometriosis is an inflammatory gynaecologic disease characterized by ectopic growth of endometrial-like tissue, resulting in pelvic pain and infertility. T-helper 9 (Th9) cells play a known role in various chronic inflammatory diseases. Despite parallels between endometriosis and Th9-driven diseases, their role in endometriosis has not been explored. We investigated Th9 cell involvement in endometriosis pathophysiology using human tissue samples, *in vitro* experiments with human-derived Th9 cells, and *in vivo* experiments to shed insight on the impact of adoptively transferred Th9 cells in our established syngeneic endometriosis mouse model. Immunohistochemistry of a tissue microarray revealed significantly increased interleukin-9 (IL-9)-positive cells in patient lesions compared to control endometrium. Human CD4+ Th cells purified from peripheral blood mononuclear cells treated with Th9-driving growth factors produced significantly increased pro-inflammatory mediators, including IL-5, IL-9 and IL-13, in response to estrogen stimulation. Adoptive transfer of murine Th9-like cells increased plasma IL-1α concentration and altered transcriptional profiles of several signalling pathways, including Notch and PI3K-Akt. Immunofluorescent microscopy depicted adoptively transferred Th9 cells present within mouse lesions. Furthermore, immunohistochemical analysis demonstrated reduced lesion proliferation following Th9-adoptive transfer. This study provides the first evidence that Th9 cells likely promote immune-inflammatory alterations within lesions to exacerbate disease.

## Introduction

Endometriosis is a chronic inflammatory disease wherein endometrial-like (endometriotic) tissue proliferates at ectopic sites, most commonly on peritoneal surfaces and pelvic organs. Our group and others have provided evidence that immune dysregulation in endometriosis is a predominant mechanism associated with disease progression (1–3). This, combined with an estrogen-dominant, progesterone-resistant endocrine imbalance further shapes the immune landscape.

T-helper type 9 (Th9) cells are a specialized Th subset named for their production of interleukin 9 (IL-9). These cells are involved in allergic responses and inflammation in parasitic infections, parallel to the classical roles of mast cells. Th9 cells differentiate from naïve CD4+ T cells when exposed to IL-4 and TGF-β, but can also differentiate from Th2 cells when stimulated with TGF-β (4, 5). While its function and regulation are still not fully understood, IL-9 is known as a pleiotropic cytokine with roles in T cell and mast cell development, and has been implicated in numerous pathologies as a regulator of inflammatory and proliferative signals (6, 7). In addition, IL-9 has been identified for its role in fibrosis of the lungs and airway in subepithelial compartments (8), and was reported as a reliable marker of poor healing in ulcerative colitis (9).

Evidence in literature indicates that IL-9 signalling is elevated in endometriosis. In 2012, Lessey *et al.* documented increased concentrations of IL-9 in the peritoneal fluid of endometriosis patients (10). We also previously demonstrated that IL-9 levels were significantly higher in plasma of patients compared to fertile healthy controls, and that endometriotic lesions as well as matched patient eutopic endometrium produced IL-9 (11). More recently, Tarumi *et al.* identified a significantly higher presence of IL-9+ CD4+ immune cells in the peritoneal fluid of endometriosis patients compared to controls (12).

The dysregulation of several Th subtype populations has been documented in endometriosis pathophysiology. Our group recently demonstrated the dysregulated functioning of Th17 cells in endometriosis (13), and the disease’s overall immune dysregulation is known to skew toward Th2 responses (14). Furthermore, we have previously reported on the involvement of mast cells in endometriosis (15), a cell type that relies on IL-9 for development. However, the involvement of Th9 cells and IL-9 in endometriosis has not yet been explored. Th9 cells have been identified to have important roles in allergic and autoimmune diseases, cystic fibrosis, and several cancers including endometrial carcinoma, where IL-9 was regulated by progesterone receptor expression (7, 16).

Inflammation and proliferation are central to endometriotic lesion progression and the alteration of these processes by immune dysregulation is continually being investigated. Here, we address the knowledge gap surrounding the potential role of Th9 cells in modulating endometriotic lesion-associated immune inflammatory alterations. Our study identifies the elevated presence of IL-9 in human endometriosis lesions, examines the effects of estradiol (E2) and progesterone (P4) on the cytokine secretory profile of Th9-driven human T cells, and evaluates the impacts of Th9 cell adoptive transfer in a mouse model of endometriosis.

## Results

### IL-9-positive cells were significantly higher in endometriosis patient lesions compared to control endometrium

To quantify IL-9 presence in endometriosis lesions, immunohistochemical staining for IL-9 was performed on a tissue microarray comprised of patient (ovarian endometriosis lesions and matched eutopic endometrium) and control endometrial samples. Image analysis with optimized HALO AI cytonuclear algorithm found significantly higher percentage of IL-9^+^ cells (p= 0.0419) in endometriosis lesions (endometrioma) compared to control endometrium (Figure 1). All control endometrium samples were in proliferative phase. Among the 12 patient eutopic endometrium samples, 5 were in secretory phase, 5 were in proliferative phase, and two were inactive endometrium. IL-9 positive cells were predominantly found within the stroma, with some glandular epithelial cells comprising the positive cell population, and a small percentage (0.2-3%) represented within the endothelial cell compartments. In lesions, stromal cells represented the majority of IL-9+ cells, being significantly higher than IL-9+ epithelial cells (p<0.0001). A similar result was observed in the inactive eutopic endometrium with p= 0.0072. However, the n= 2 for inactive endometrium samples was insufficient to produce representative statistical results. Meanwhile, no significant difference was observed between stromal and epithelial IL-9+ cells for control endometrium or secretory or proliferative eutopic endometrium. Further, the proportion of IL-9+ cells represented by stromal cells in lesions was significantly higher than that of all other sample types, excluding inactive endometrium (Supplemental Figure 1).

**Figure 1.**
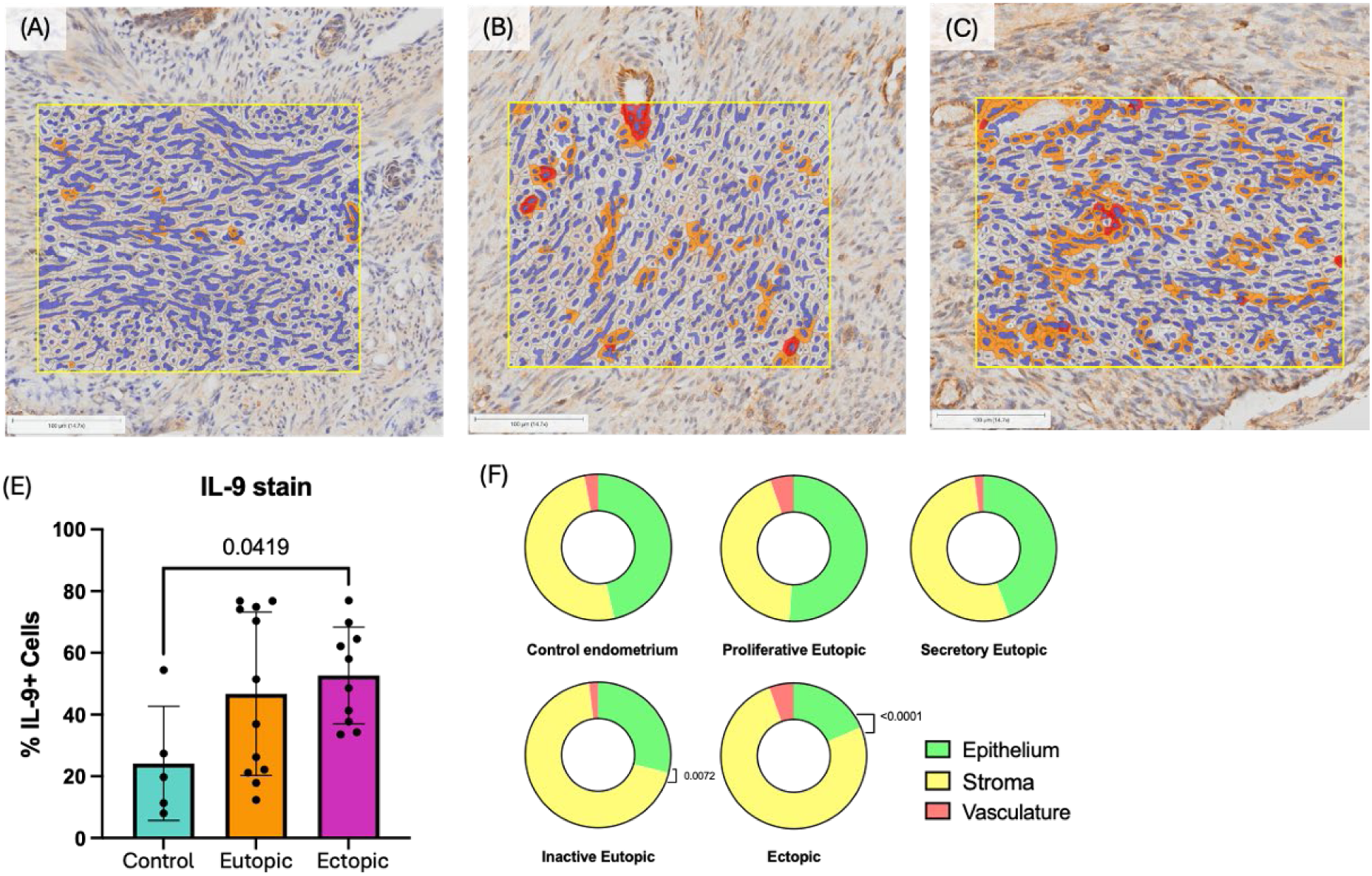
IL-9 presence in human endometrioma, eutopic endometrium and control endometrium. Eighty-one cores in tissue microarray containing triplicates of control endometrium (n=5), eutopic patient endometrium (n=12) and endometrioma tissues (n=10) were stained with anti-IL-9 antibody. HALO AI (Indica Labs) was used to classify tissue into classes of epithelium, stroma, and vasculature, with classes to exclude glandular lumen, slide glass, and debris from analysis. Cytonuclear analysis algorithm was optimized to detect weak (yellow), moderate (orange) and strong (red) anti-IL-9 stain. (A) Endometrioma lesion with cytonuclear analysis markup showing positive IL-9 staining. (B) Eutopic endometrium from endometriosis patient with cytonuclear analysis markup showing positive IL-9 staining. (C) Control healthy endometrium with cytonuclear analysis markup showing sparse staining for IL-9. (E) Percent cells positively stained for IL-9 in control endometrium, eutopic endometrium, and endometrioma lesion tissues. Staining of IL-9 in endometrioma tissues is significantly stronger than control endometrium (p= 0.0419). (F) Proportion of IL-9+ stromal cells was strongest in ectopic samples, significantly higher than epithelial proportion (p>0.0001). This difference was not observed in control, proliferative eutopic, or secretory eutopic endometrium. Inactive endometrium (n= 2) showed a similar disparity in IL-9+ representation between stromal and epithelial cells.

### IL-7 increased secretion of pro-inflammatory, chemotactic, and angiogenic cytokines by Th9-driven CD4+ human lymphocytes

Using human peripheral blood mononuclear cells (PBMCs) isolated from healthy volunteer blood, CD4+ T cells were isolated by immunomagnetic negative selection and cultured with Th9 growth factors (IL-4 and TGF-β) to drive them toward a Th9 phenotype. With the aim of optimizing Th9 cell differentiation, IL-7 was included in the Th9-driving cocktail based on literature suggesting that IL-7 increases IL-9 production by Th9 differentiated cells *in vitro* (17). Indeed, groups that received IL-7 had significantly increased concentration of IL-2, IL-8, IL-17A, sCD40L, GM-CSF, and TNF-α (Figure 2A—F).

**Figure 2.**
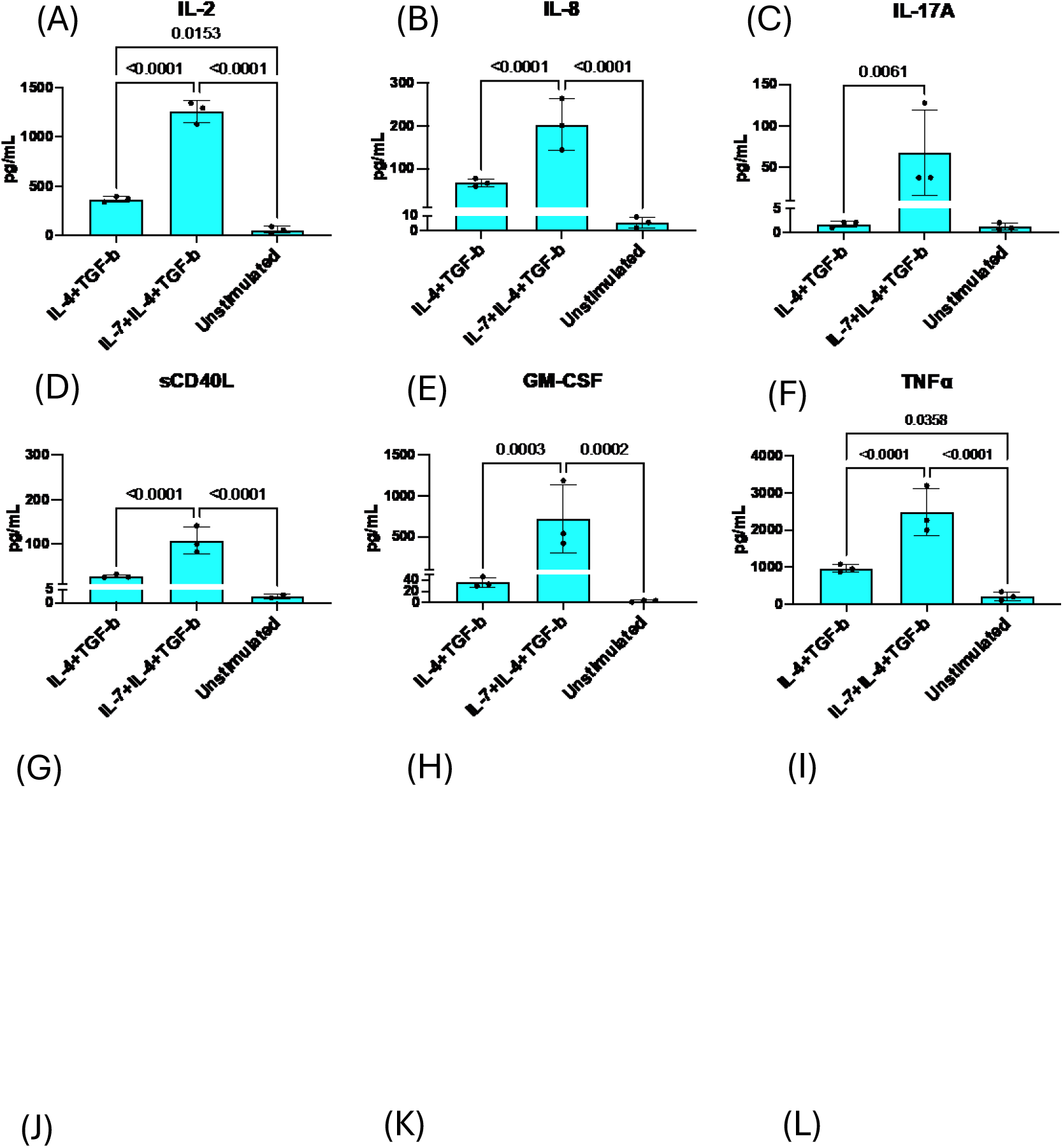
Cytokine response by PBMC-derived Th9-driven T cells to different growth factor cocktails. CD4+ PBMCs were negatively selected by immunomagnetic separation and cultured in media conditions to drive Th9 phenotype development (IL-2, IL-4, TGF-β, anti-CD3ε, anti-CD28, anti-IFNγ) with or without IL-7, or without any growth factor cocktail (unstimulated). Cultures were restimulated with PMA and ionomycin for 5 hours before supernatant was collected and analyzed with a multiplex cytokine analysis panel of 32 inflammatory cytokines (Eve Technologies, Calgary, AB). As shown in A–F, the production levels of several cytokines by PBMC-derived Th9-driven T cells were significantly increased by the inclusion of IL-7 in media. G–L: Culture conditions were as described, with hormonal treatment groups including 17-β-estradiol [1x10^-7^ M] (E2) and progesterone [1x10^-7^ M] (P4). As shown in G–L, the production levels of several cytokines by PBMC-derived Th9-driven T cells were significantly increased by the inclusion of E2 treatment. Notably, classical Th2 cytokines IL-5, IL-9, and IL-13 were significantly increased by E2.

### Estrogen increased IL-9 secretion in Th9-driven CD4+ human lymphocytes

To evaluate the impact of E2 and P4 hormones on the secretory profile of human PBMC-derived Th9-driven T cells, groups were treated with growth factors as detailed above and treated 24-hours with 1.0x10^-6^ M E2 or 1.0x10^-6^ M P4. In the supernatant of PBMC-derived Th9-driven T cell cultures, multiplex cytokine analysis revealed that cells treated with E2 had significantly increased concentration of IL-9, IL-5, IL-13, IL-17F, CCL22, and CXCL9 compared to unstimulated Th0 cells and cells treated with the Th9-driving cocktail but not E2 or P4 (Figure 2G—L). These findings suggest estrogen may have an impact on the secretory profile of *in vitro* derived human Th9 cells.

### IL-9R+ peritoneal immune cells were increased in endometriotic mice compared to sham-operated controls

Flow cytometric analysis was used to evaluate the presence of IL-9R (IL-9 receptor) expression in peritoneal immune cells of mice induced with endometriosis vs. sham operated controls. Compared to sham operated mice, endometriosis-induced mice showed significantly higher presence of IL-9R^+^ cells within the CD45^+^ peritoneal cell population (p= 0.031, Figure 3A). As multiple pathogenic mechanisms of Th9 cells have been reported to depend upon IL-9 stimulation of mast cells (MCs)(18, 19), we also evaluated IL-9R expression in both MCs and committed mast cell progenitors (MCcp). No significant difference was observed in frequency of MCs or Th9 cells in peritoneal fluid or splenocyte populations between endometriosis-induced and sham operated mice (data not shown). No difference in MC IL-9R expression was found between the groups (Figure 3B). However, within the MCcp populations, expression of IL-9R was significantly lower in MCcp of mice induced with endometriosis compared to sham operated mice (p= 0.034, Figure 3C). While the overall MCcp population was significantly larger within peritoneal immune cells of sham-operated controls compared to endometriosis-induced mice (p= 0.00441, data not shown), this stark difference in IL-9R expression is noteworthy considering IL-9R expression in CD45+ peritoneal cells was higher in endometriosis-induced mice.

**Figure 3.**
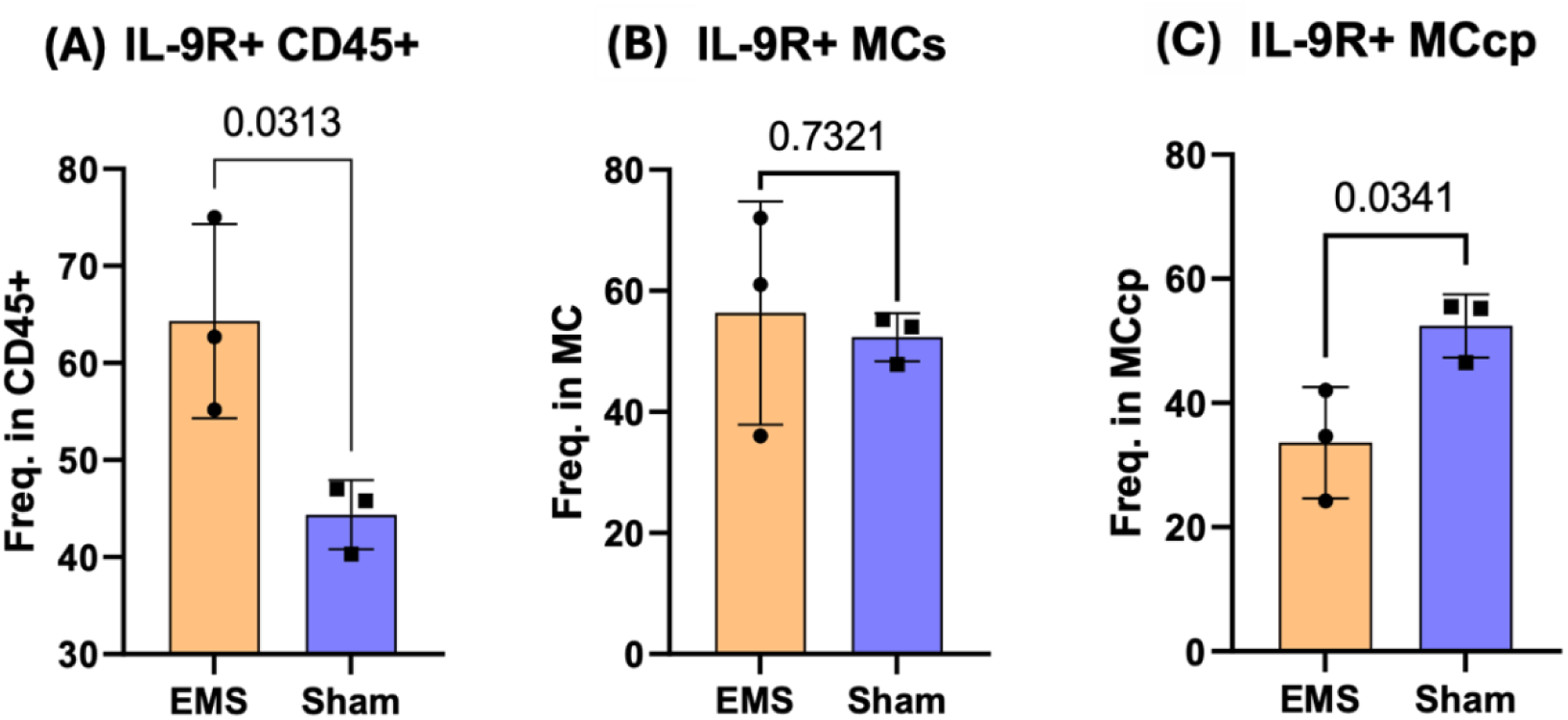
IL-9R^+^ peritoneal immune cell populations of mice induced with endometriosis or sham-operated controls. (A) IL-9R+ cells were significantly higher in CD45+ peritoneal cells of mice induced with endometriosis compared to sham-operated controls. (B) No significant difference in IL-9R expression within mast cells in endometriosis-induced mice vs. sham-operated controls. (C) Out of the committed mast cell progenitor populations in each group, a significantly higher portion were IL-9R+ in the sham-operated group compared to the endometriosis-induced group. “EMS”= endometriosis-induced mice; “Sham”= sham-operated control mice.

### In vitro derivation of Th9 cells from mouse splenocytes

To gain insights into the influence of adoptively transferred Th9 cells on the immune microenvironment of murine endometriotic lesions, we first aimed to derive Th9 cells *in vitro* from murine splenocytes. Using an optimized format of Pham’s protocol (20) to derive Th9 cells, the Th9 growth cocktail-treated group yielded significantly higher Th9 cells compared to those without Th9 growth factors as well as the monensin-treated group (p≤0.05, Figure 4B). Monensin is a protein transport inhibitor used to prevent secretion of soluble cytokines and allow for intracellular staining, and its reduction of Th9 cell yield likely occurred because Th9 cells require autocrine IL-2 and TGF-β stimulation for phenotype development (20). Treating the Th9-driven group with E2 [1x10^-6^] had no significant impact on Th9 yield, but showed a slight downward trend compared to the group treated only with Th9 growth factors. The confirmation of mouse Th9 cell fate and expansion *in vitro* carried forward subsequent adoptive transfer experiments in our mouse model of endometriosis. Cells that were CD8α ^—^/CD14^—^/CD19^—^/CD4^+^/IRF4^+^/ IL-4Rα^+^ were considered Th9 cells (Figure 4A).

**Figure 4.**
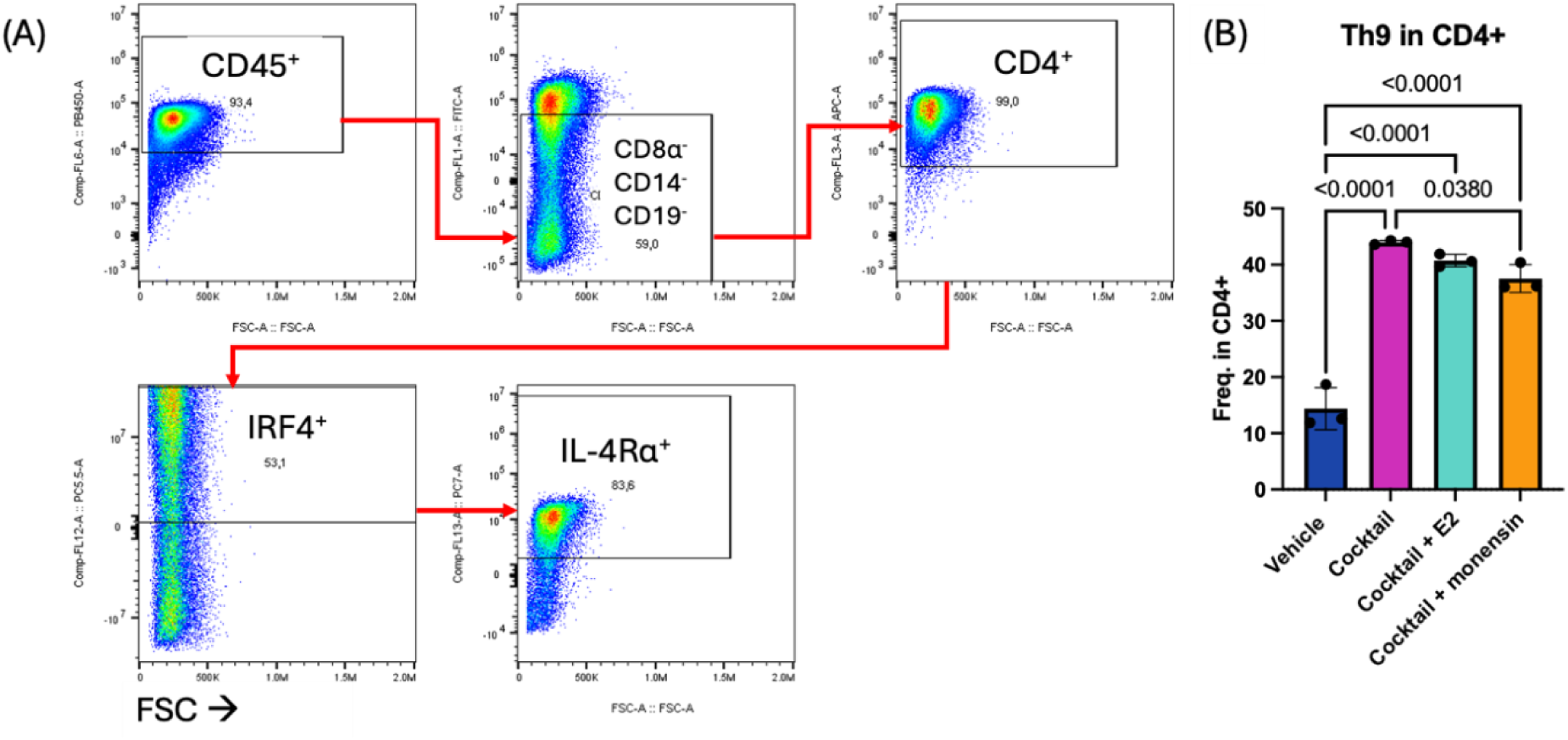
Deriving Th9 phenotype *in vitro* from mouse splenocytes. CD4+ cells were negatively selected using immunomagnetic separation and cultured with a cocktail of growth factors (IL-2, IL-4, TGF-β, anti-CD3, anti-CD28, anti-IFNγ). Cells were stimulated with PMA and ionomycin for 6 hours. (A) Gating strategy for selecting Th9 phenotype. Out of CD45^+^ population, cells negative for CD8α, CD14, and CD19 were selected. The CD4^+^IRF4^+^IL-4Rα^+^ cells within this CD45^+^CD8a^-^CD14^-^CD19^-^ population were considered Th9 cells. (B) One-way ANOVA test showed using the growth cocktail yielded significantly higher cells of the Th9 phenotype than vehicle control. Use of monensin significantly decreased phenotypic yield compared to growth cocktail without monensin.

### IL-1α significantly increased in plasma following Th9 adoptive transfer in a mouse model of endometriosis

After murine induction of endometriosis on day 0, blood was collected at day 7 and 14 via submandibular vein puncture to measure plasma cytokine levels before and after adoptive transfer of Th9-like cells. Analysis revealed that IL-1α was significantly increased (p= 0.0276) between day 7 and day 14 in the Th9-like adoptive transfer group, while no changes were observed between these time points for the PBS control group (Figure 5B). This suggests that adoptively transferred Th9 cells were likely exerting a pro-inflammatory response captured in the systemic circulation.

**Figure 5.**
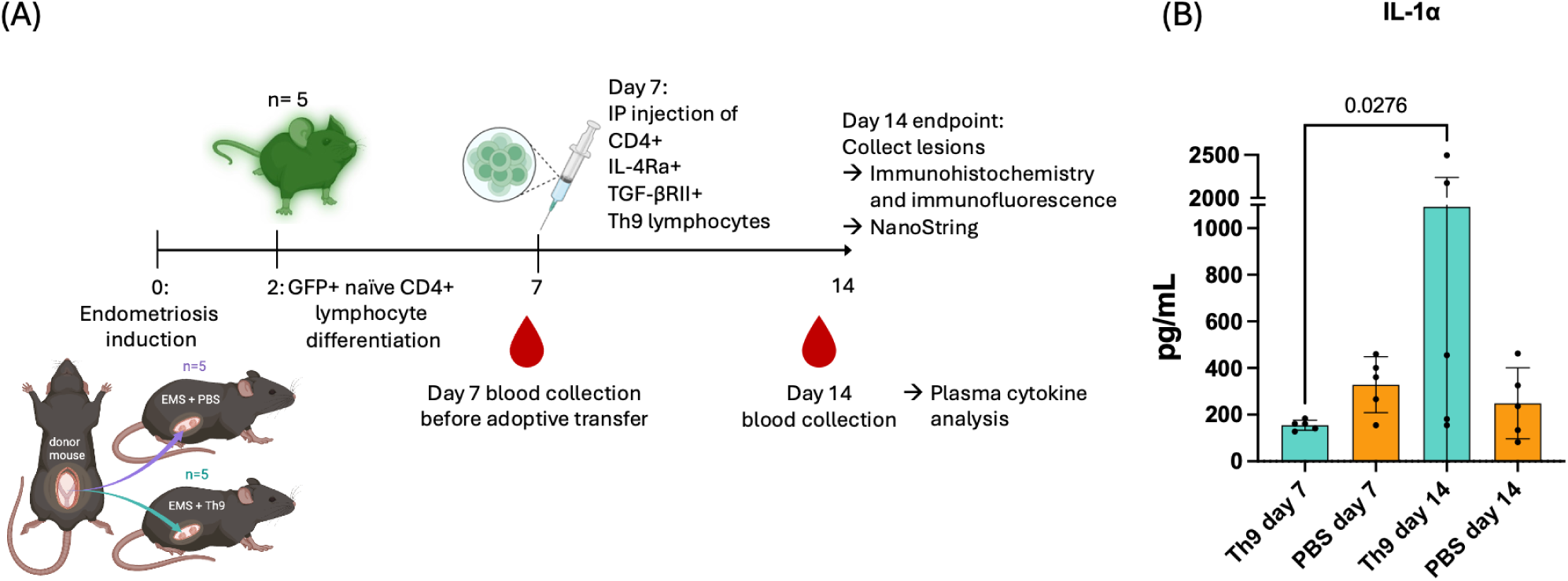
Endometriosis mouse model with adoptive transfer of GFP+ Th9 cells or PBS. (A) Workflow of endometriosis mouse model with adoptive transfer of Th9-like cells. Mice were induced with endometriosis on day 0. GFP+ mouse splenic CD4+ T cells were cultured *in vitro* with Th9 phenotype-deriving growth factors and double positively selected for IL-4Rα and TGF-βRII^+^ by immunomagnetic separation. On day 7, mice were injected intraperitoneally with suspensions of 2.74x10^5^ Th9-like lymphocytes or PBS. (B) Mouse plasma collected at 7- and 14- day timepoints underwent multiplex analysis for inflammatory cytokine levels. IL-1α significantly increased in the Th9 adoptive transfer group from day 7 to day 14 while no change was observed in the PBS control group. “Th9” = mouse group that received adoptive transfer of Th9-like lymphocytes; “PBS”= PBS control mouse group.

### Th9 cells infiltrated endometriotic lesions upon adoptive transfer in a mouse model of endometriosis

Following adoptive transfer of GFP+ Th9-like lymphocytes into mice induced with endometriosis, we aimed to establish whether these adoptively transferred cells infiltrated into endometriotic lesions. Indeed, we identified GPF+ CD4+ IL-9+ cells incorporated into endometriotic lesions through fluorescent microscopy of mouse lesion tissue (Figure 6). GFP+ cells were evenly distributed throughout stromal compartments, with very few cells localized to epithelial compartments. This qualitative observation supports the notion that adoptively transferred Th9-like cells likely contribute to lesion pathology and associated microenvironmental changes.

**Figure 6.**
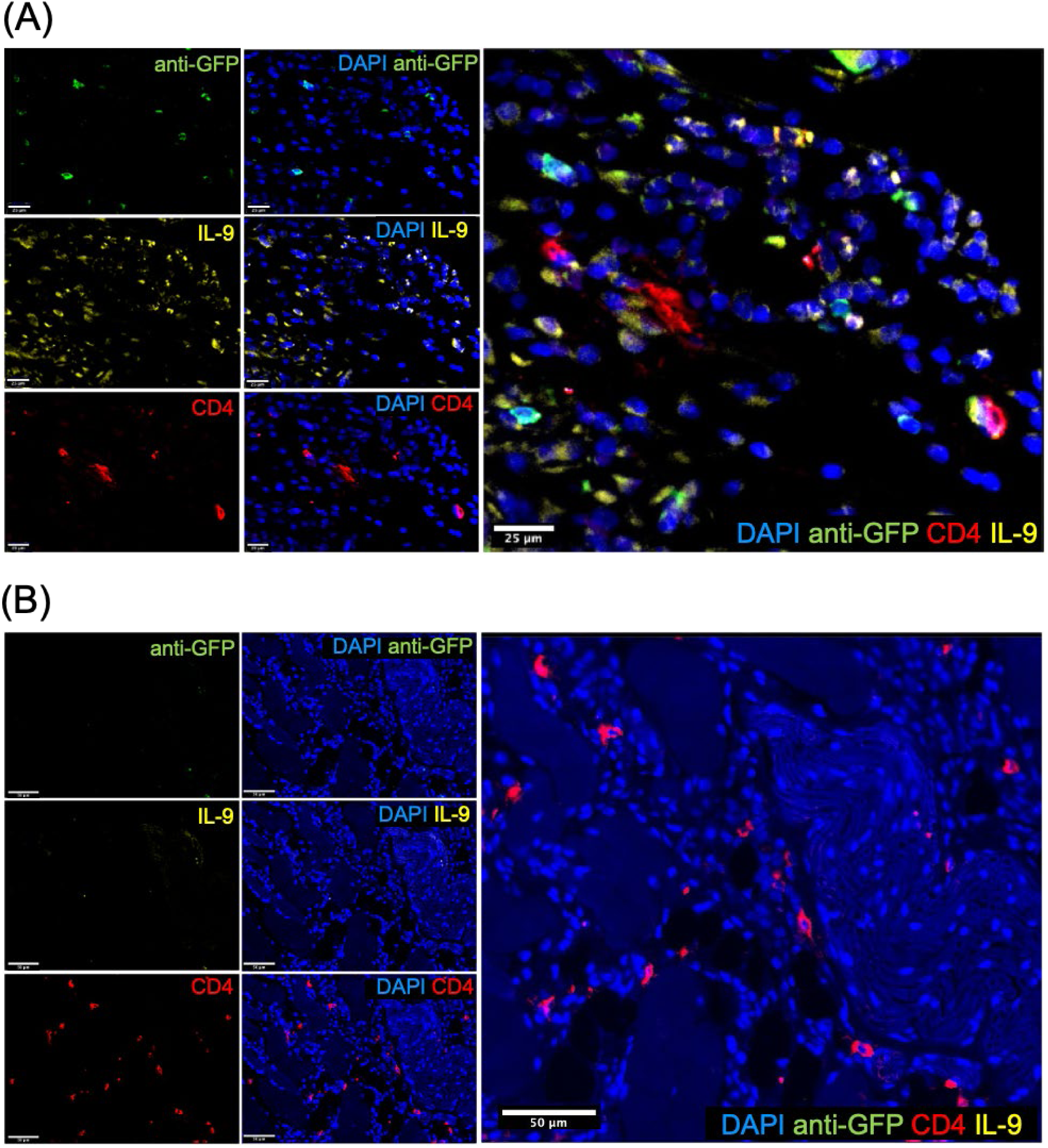
Adoptively transferred GFP+ Th9-like cells infiltrated into mouse endometriosis lesion tissue. GFP+ CD4+ IL-9+ cells observed in lesions of mice that received Th9 cells (A). Control mouse lesion displayed some CD4+ and IL-9+ staining but lacked GFP+ signal (B).Blue= DAPI nuclear stain, green= anti-GFP—Alexa Fluor 488, yellow= anti-IL-9—Alexa Fluor 568, red= anti-CD4—Alexa Fluor 647.

### Mouse endometriotic lesion proliferation was significantly impeded following Th9 adoptive transfer

As proliferation is a hallmark of endometriosis lesion establishment and subsequent progression, and Th9 cells have been reported to affect proliferation in various cell types, we performed immunohistochemistry on mouse lesion tissues for Ki67, a proliferation marker. Analysis revealed significant reduction (p= 0.0059) of Ki67 immunostaining (mean percentage of Ki67-positive cells) in the lesions of mice that received adoptive transfer of Th9-like lymphocytes as compared to the PBS control group (Figure 7).

**Figure 7.**
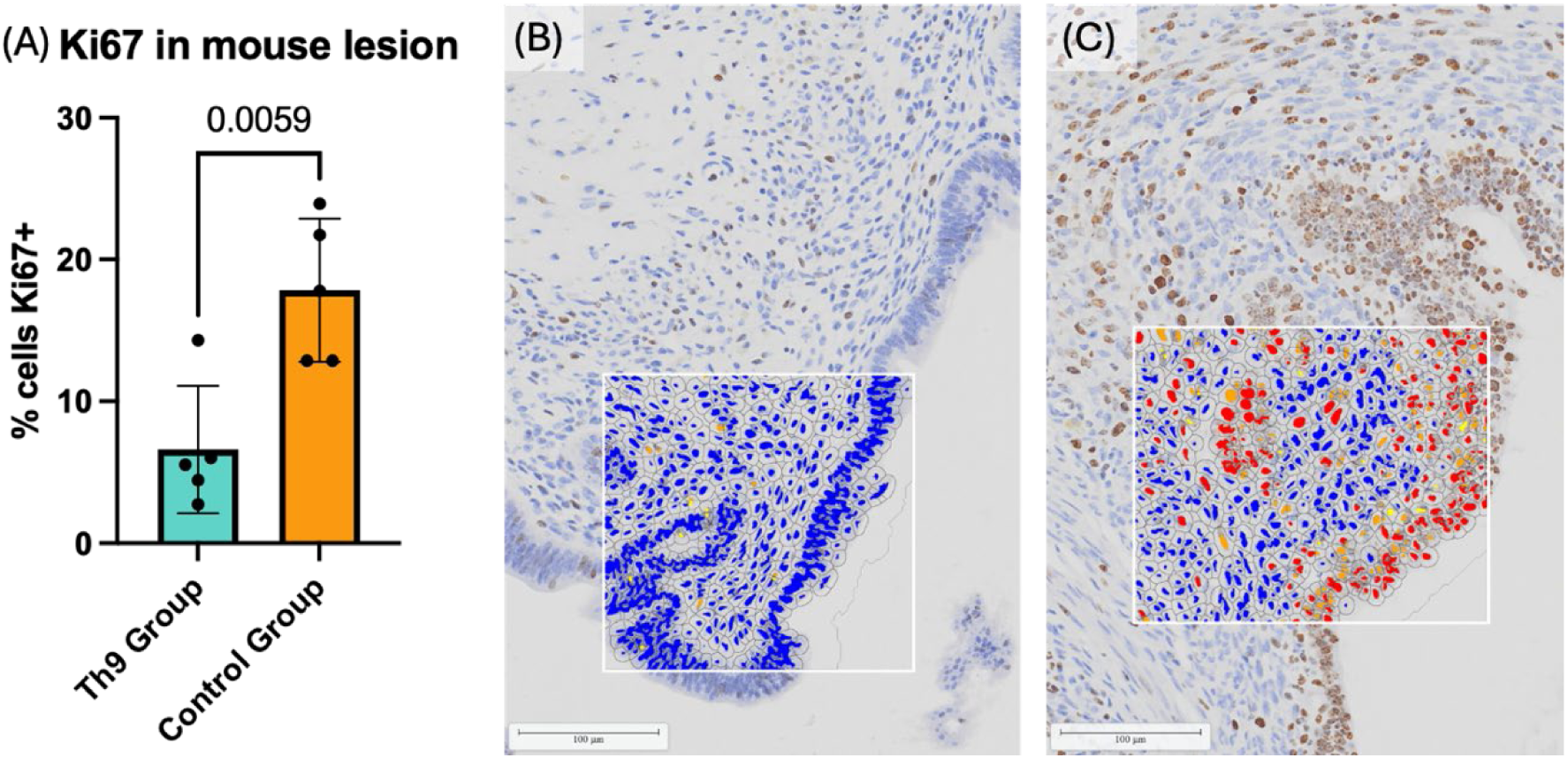
Ki67 immunohistochemical staining in endometriotic lesions of mice who received adoptive transfer of Th9 cells or PBS control. (A) Percentage of Ki67+ cells was significantly higher in the PBS control group (mean of 17.8% Ki67+ cells) compared to the Th9 adoptive transfer group (mean of 6.6% Ki67+ cells). (B) Endometriotic lesion tissue from a mouse that received adoptive transfer of Th9-like cells. Ki67 positive staining is sparse and weak. (C) Endometriotic lesion tissue from a mouse that received PBS control. Ki67 positive staining is strong, indicating proliferative profile typical of endometriotic lesions in this model. Scale bars= 100µm. Blue= nucleus, yellow= weak positive stain, orange= moderate positive stain, red= strong positive stain.

### Adoptive transfer of Th9 cells altered the transcriptional profile of murine endometriotic lesions

To further understand the impact of adoptively transferred Th9-like cells in murine lesion microenvironment, total RNA was extracted from lesions and evaluated with the nCounter Mouse Fibrosis V2 panel from NanoString. This panel includes a broad list of genes involved in inflammation, immunity and fibrosis. Analysis of data in nSolver software revealed 43 genes were significantly differentially expressed (p≤0.05). Among them, 39 genes were significantly upregulated in the Th9 adoptive transfer group and four genes (*Banf1*, *Hsp90ab1*, *Prkag2*, *Traf6*) were significantly downregulated (Figure 8). Samples were analyzed with unsupervised clustering, but clustered almost exclusively within their respective treatment groups because of their similarities in gene expression profiles. Out of the 39 genes significantly upregulated in lesions of the Th9 adoptive transfer group, 20 genes were associated with the process of proliferation and 15 were associated with inflammation. Several pathways and themes are represented in these results; three genes associated with extracellular matrix (ECM) formation (*Col5a3*, *Itga4*, and *Mfap3*), six genes relevant in Notch signalling (*Adam17, Jag1, Notch4, Ppard, Psenen*, *Psen2*), and seven genes associated with the adenosine pathway (*Adcy7*, *Arhgef2*, *Arrb1*, *Pde2a*, *Plcb2*, *Ptger4*, *Sdc3*) were significantly upregulated in the lesions of the Th9 adoptive transfer group. Five genes involved in PI3K-Akt signalling (*Pi3kr5*, *Akt1*, *Phlpp1*, *Itga4*, *Creb3*) were significantly upregulated while another PI3K-Akt-associated gene, *Hsp90ab1,* was downregulated in the Th9 cell adoptive transfer group.

**Figure 8.**
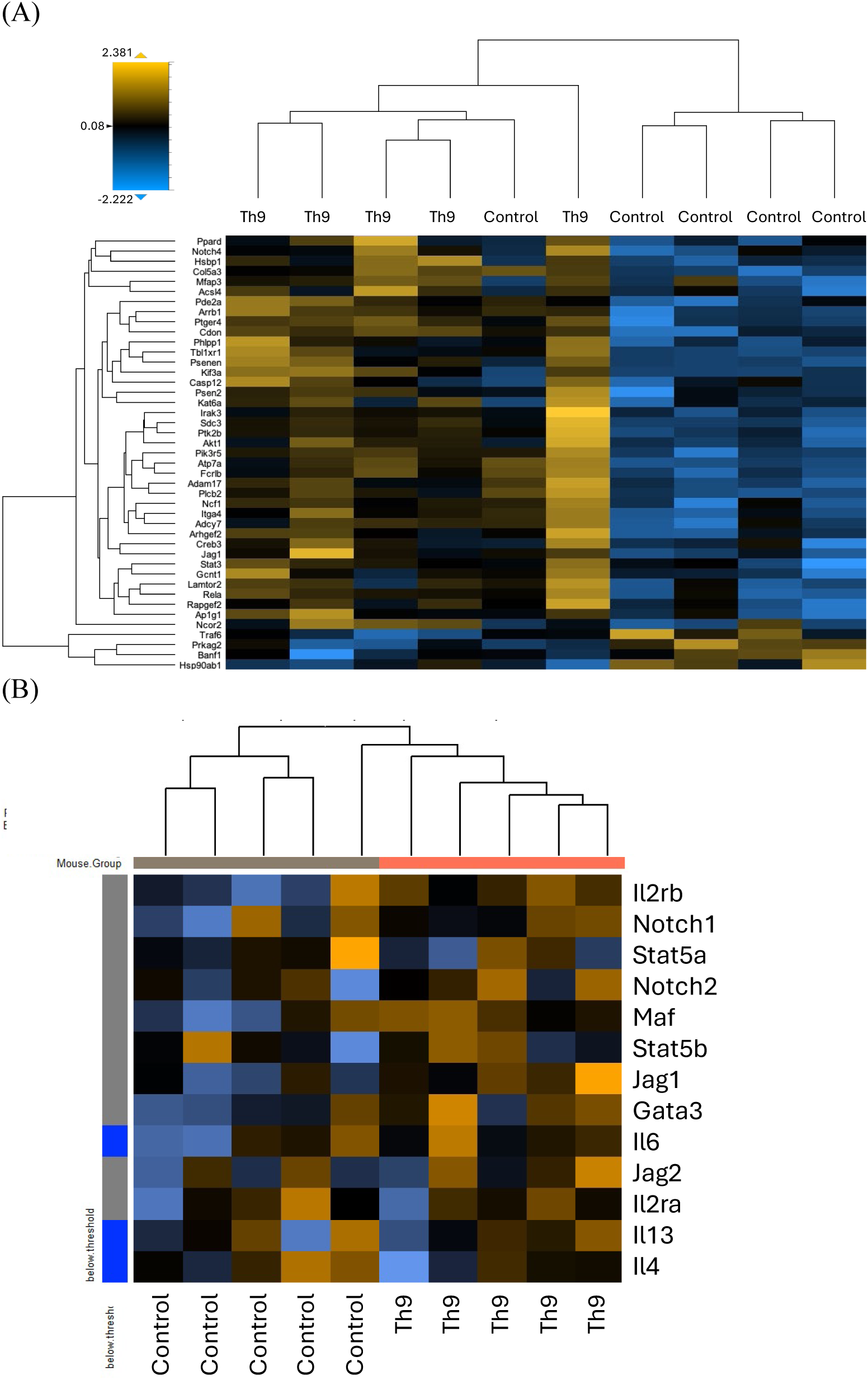
Heat map of significant differentially expressed genes in mouse endometriotic lesions of mice that received Th9 adoptive transfer or PBS. RNA samples isolated from endometriotic lesion tissues of mice that received adoptive transfer of Th9-like cells (n= 5) or PBS (n= 5) were evaluated for expression of 760 genes using NanoString’s nCounter Mouse Fibrosis V2 panel. Samples were analyzed with unsupervised clustering. Dendogram across the top of the heat map shows clustering of samples almost exclusively into separate groups. (A) Thirty-nine genes were significantly upregulated in the Th9 adoptive transfer group while 4 were significantly downregulated. (B) Using unsupervised clustering, samples clustered together with their respective treatment groups. Genes involved in Th2 differentiation were generally upregulated in the Th9 adoptive transfer group compared to the control group. Orange= increased expression, blue= decreased expression

### Th2 differentiation transcripts were altered in lesions of mice who received Th9 cell adoptive transfer

Advanced analysis in nSolver software was conducted to identify consistent trends in gene expression. Genes associated with Th2 differentiation (*Gata3*, *Il13*, *Il2ra*, *Il2rb*, *Il4*, *Il6*, *Jag1*, *Jag2*, *Maf, Notch1*, *Notch2*, *Stat5a*, *Stat5b*) were found to be upregulated in the Th9-adoptive transfer group, with *Jag1* significantly increased (Figure 9). *Stat3*, though not included in this heat map, is required for Th2 development and was significantly upregulated in the Th9 adoptive transfer group. This preliminary analysis further supports the influence of Th9 cells on the Th2 inflammatory response.

### Gene pathways for focal adhesion, PI3K-Akt, chemokine signalling, and cytokine-cytokine receptor interaction were altered in endometriotic lesions of Th9 cell adoptive transfer group

Within the nSolver advanced analysis, Pathview analysis highlighted a significant difference in genes involved in focal adhesion, wherein the expression levels of several genes were lower in the PBS control group as compared to the Th9 adoptive transfer group. Specifically, genes within the integrin α (ITGA) family, receptor tyrosine kinase (RTK), with downstream effectors including Src, Crk, and Rac.

## Discussion

Endometriosis is a complex inflammatory condition with a well-documented bias for Th2 immune responses (14, 21, 22). As Th9 cells regulate Th2 response by secreting IL-9, it is possible that Th9 cells are involved in endometriosis pathophysiology. However, there have been no reports thus far documenting the presence or involvement of Th9 cells in endometriosis. Our group has extensively reported on various pathways driving immune dysregulation in endometriosis, including the involvement of estrogen-mediated mast cells and the dysregulated pathogenic IL-23/Th17 axis in endometriosis(13, 15). Considering evidence that Th9 cells cooperate with mast cells and other immune cells to progress various inflammatory diseases (7), we investigated the potential role of Th9 cells in endometriosis through *in vitro* derivation of this phenotype and adoptive transfer of Th9-like lymphocytes in mouse model of endometriosis.

The observed increase in IL-9 protein in human endometriotic lesions is congruent with literature depicting increased IL-9 in peritoneal fluid, as well as our previous findings of increased IL-9 levels in endometriosis patient plasma (11, 23). Further, the proportion of stromal cells within IL-9+ cells was significantly higher in lesion samples, illustrating a distinction in IL-9 distribution in the lesion microenvironment as compared to normal or eutopic endometrium. Given inactive endometrium is marked by an increase in fibrosis(24), the potential relationship of IL-9 in fibrosis of endometriosis lesions should be investigated further. As discussed, IL-9 is a growth factor for mast cells and other immune cells that can promote inflammation by supporting mast cell development (25). Though, other inflammatory pathways, particularly in relation to type 2 inflammation, are also relevant; in 2011, Blom *et al.* identified that human CD4+ T cells can be induced by IL-33 to secrete IL-9 (22). Conversely, Du *et al.* found that IL-9 was a downstream effector of IL-33 in airway inflammation, and that IL-9 deficient mice experienced significantly decreased effects of their experimental IL-33 challenge (27). Our group has established the role of IL-33 in activating group 2 innate lymphoid cells in endometriosis progression (28). The inflammatory relationship between IL-9 and IL-33 has yet to be explored in endometriosis pathophysiology, and will be a worthy area of future investigation. While our finding of IL-9 presence is significant, the lesion tissues in this study only represented the ovarian (endometrioma) phenotype of pelvic endometriosis. As this disease is not monolithic, there remains a need to investigate the presence and involvement of IL-9 in all phenotypes and stages of endometriosis.

To gain further insights into the complex biology of Th9 cells, CD4+ T cells were isolated from human PBMCs and cultured in the presence of IL-7, IL-4, and TGF-β to drive a Th9 phenotype. *In vitro* treatment of E2 and P4 was examined to understand hormonal influences on Th9 cells. Cytokine profiling revealed that inclusion of IL-7 led to significant increase of IL-2, IL-8, IL-17A, sCD40L, and TNF-α (Figure 2A—F). Overall, these results can be expected as IL-7 is known to support T cell development and homeostasis (29). Further, stimulation with IL-7 in combination with E2 significantly increased IL-9 production by human PBMC-derived Th9-driven T cells. This suggests E2 may influence IL-9 production by Th9-like cells. (30)(10)

Ultimately, the dysregulation of IL-9 signalling in endometriosis is part of a complex network of dysregulated immune pathways. In 2015, our group showed that Ishikawa cells (endometrial adenocarcinoma cell line analog of endometrial stromal cells) treated with IL-17A significantly increased production of IL-9 (30). We subsequently reported in 2016 that endometriosis patient plasma contained significantly higher levels of IL-9 as compared to control plasma (11). We have since expanded upon the role of IL-17 and Th17 cells in endometriosis, while other groups have reported on the estrogenic regulation of IL-17 production (31). Our *in vitro* treatment of PBMC-derived Th9-driven lymphocytes aimed to understand the effects of estrogen on the secretory profile of Th9 cells. Multiple research groups have demonstrated the importance of estrogen and estrogen receptor α in T cell functioning and activation (32, 33). Indeed, in our *in vitro* experiments, PBMC-derived Th9-driven T cells responded to estrogen stimulation with increased production of IL-5, IL-9, IL-13, IL-17F, CCL22 and CXCL9. This profile appears partially in line with promoting type 2 inflammation, along with chemotactic signalling which would serve to recruit lymphocytes and augment the local immune microenvironment. We fully recognize the complex interactions between various immune cells, cytokines and growth factors in shaping the chronic nature of endometriosis-associated inflammation, as discussed in several of our previous review articles (14, 34, 35).

Since several of the documented pathogenic roles of Th9 cells are found to be exerted through their support of mast cells with IL-9 (18, 36), we used flow cytometry to analyze IL-9R expression in mast cells from the peritoneal fluid of mice induced with endometriosis. While IL-9R expression in mast cells did not significantly differ between endometriosis and sham-operated mice, expression of IL-9R in CD45+ peritoneal cells was significantly increased in endometriotic mice compared to sham-operated controls. This is relevant to findings from Tarumi *et al.* in human patient tissue, where IL-9R immunostaining was significantly higher in ovarian endometriotic lesions compared to eutopic endometrium and normal endometrium(12). It will be important to further examine which cell types comprise this IL-9R+ population to understand how IL-9 signalling impacts the immune microenvironment of endometriosis.

In endometriosis-induced mice that received adoptive transfer of GFP+ Th9 cells, we detected GFP+ CD4+ IL-9+ cells in mouse endometriotic lesions. In addition to verifying the migration of these cells into the lesion tissue, we captured significant impacts on the proliferation and transcriptional prolife of endometriotic lesions. Specifically, adoptive transfer of Th9-like lymphocytes had anti-proliferative effects in mouse endometriotic lesions. While immunohistochemical staining for Ki67 in mouse lesion tissue showed this contrast quantitatively, more intricate details of this effect were unearthed in the analysis of differently expressed genes (DEGs) in RNA from mouse lesions. Of the 39 genes found to be significantly upregulated in lesions of the Th9 adoptive transfer group, 20 genes were associated with the process of proliferation and 15 were associated with inflammation.

While few of the genes examined for Th2 differentiation were significantly differentially expressed, advanced analysis in nSolver software detected a general upward trend of this transcriptional profile in the Th9 adoptive transfer group. Given the plasticity and overlap between Th2 and Th9 biology, this overall trend might be explained by the adoptive transfer of a high number of Th9-like cells. This is especially plausible considering the Th9 adoptive transfer group saw significantly increased expression of *Stat3*, a critical driving factor in Th2 differentiation that is also activated in Th9 differentiation (4).

Six genes relevant in Notch signalling (*Adam17, Jag1, Notch4*, *Ppard*, *Psenen*, *Psen2*) were significantly increased in the Th9 adoptive transfer group. Notch signalling is required in Th2 differentiation and its role in the regulation of Th9 biology has been identified. Specifically, research shows ablating Notch1 and Notch2 receptors inhibits Th9 development, while Notch pathway activation downstream of TGF-β stimulation induces IL-9 secretion independent of IL-4 (37). *Psenen* and *Psen2* are integral to the γ-secretase complex, which processes Notch receptors for the release Notch intracellular domains (38). Notably, Jiang *et al.* found that inhibiting Notch signalling with γ-secretase inhibitor DAPT led to inhibition of endometriosis progression in their mouse model, an effect that correlated with a reduction in myeloid-derived suppressor cells within mouse peritoneal fluid and reduced expression of *Adam17* and *Jag1* within lesion tissue (39). Metalloprotease *Adam17* plays a role in the shedding of cell surface proteins, including cytokines and their receptors, and this gene has been found to participate in regulation of endometriotic cell migration (40). Additionally, three genes associated with extracellular matrix (ECM) formation, *Col5a3*, *Itga4*, and *Mfap3*, were significantly increased in the Th9 adoptive transfer group. As discussed earlier, Th9 cells have been identified to promote fibrosis in other pathologies such as allergic airway inflammation (41).

Finally, several results of this Th9 adoptive transfer experiment indicated activated IL-1 pathway signalling. *Irak3*, the gene for IL-1 receptor associated kinase 3, was upregulated in lesions from the Th9 adoptive transfer group. From Pathview analysis in nSolver, we found multiple components of the IL-1 signalling pathway were increased in the Th9 adoptive transfer group; IL1B (IL-1β), IL1R1 (IL-1 receptor type 1), and IL1RAP (IL-1 receptor accessory protein). IL-1β stimulation has been identified as a pivotal determinant of exhaustion resistance in Th9 populations in melanoma tumours and has also been well documented for its elevated presence in endometriosis lesions (42, 43). In terms of systemic inflammatory status in the mouse model, we observed an increase in plasma IL-1α concentration in mice that received an adoptive transfer of Th9-like cells. In humans, IL-1α is also significantly increased in patient peritoneal fluid and associated with advanced disease, while its receptor antagonist, IL-1Ra, is increased during earlier stages of endometriosis (44). Aside from macrophages, monocytes and mast cells, IL-1Ra is also produced by endometrial epithelial cells (45, 46). In more advanced stages of endometriosis, inflammatory and proliferative activity have been found to wane in lesions and over time these become comprised of more stiff, fibrotic tissue (47). The involvement of Th9 cells in endometriosis could potentially be associated with later stages wherein IL-1α is increased and fibrosis dominates the lesion landscape.

The participation of Th9 cells in endometriosis may depend upon the stage and phenotype of disease. As this cell type has diverse, context-dependent roles, Th9 cells may contribute anti-proliferative functions in later stages but may facilitate mast cell-derived inflammation in earlier stages. Overall, this work has presented novel insights into the function of Th9 cells in endometriosis, their relationship with mast cells, and their impact on endometriotic lesion proliferation.

## Methods

### Study approval

Human lesion samples were obtained from patients who underwent laparoscopic excision surgery after written informed consent at Kingston General Hospital, Kingston, ON, Canada. The endometrial samples, both from patients and healthy individuals matched for uterine cycle phase were obtained by Pipelle sampling, as per standard procedure. All control endometrium samples were in proliferative phase. Among the 12 patient eutopic endometrium samples, 5 were in secretory phase, 5 were in proliferative phase, and two were inactive endometrium. Blood was collected from volunteers with informed consent by a certified phlebotomist at Queen’s Department of Biomedical Sciences, Kingston, ON, Canada. Study is approved by the Queen’s University Health Sciences Research Ethics Board, Kingston, Ontario, Canada. *In vivo* experiments were done using C57BL/6 mice acquired from Charles River Laboratories and Jackson Laboratories. Experiments were approved by the Queen’s Institutional Animal Care Committee, Kingston, Ontario, Canada.

### Evaluating IL-9 presence in tissue microarray of endometrioma lesions, eutopic endometrium, and normal control endometrium

Immunohistochemistry for anti-IL-9 antibody staining was performed on a human patient tissue microarray (TMA) of endometrioma lesions, eutopic endometrium, and control endometrium. Triplicates of control endometrium (n= 5), eutopic patient endometrium (n= 12) and endometrioma tissues (n= 10) were stained with anti-IL-9 monoclonal antibody (catalog # 66144-1-IG, Thermo Fisher Scientific). The TMA was stained using a Leica Bond RX automated stainer (Leica Microsystems). EDTA-based epitope retrieval was conducted for 10 minutes and primary antibody was incubated for 15 minutes at a concentration of 1:200. Leica BOND Polymer Refine Detection kit (Cat. DS9800, Leica Biosystems) was used for detection of 3,3’-diaminobenzidine (DAB) chromogen against a hematoxylin counterstain (Leica Microsystems). Slides were scanned at 40X magnification in an Olympus VS120 high resolution slide scanner (Olympus Life Science). Image analysis was performed using HALO AI software (Indica Labs). Scanned images were classified into slide glass (empty area), glandular epithelium, glandular lumen, stroma (proliferative and secretory), and vasculature using HALO’s tissue classifier tool in order to exclude empty areas from analysis. A cytonuclear analysis algorithm was optimized to detect weak, moderate, and strong positive staining of anti-IL-9 antibody. A one-way ANOVA test of grouped data in triplicates was used to compare percentage of positively stained cells out of all detected cells in endometrioma tissues, eutopic endometrium, and control endometrium using. Outlier test (ROUT Q= 0.5%) found zero outliers.

### Driving Th9 cells from human peripheral blood mononuclear cells

Human blood was collected by venipuncture and PBMCs were separated by centrifugation using Lymphoprep™ density gradient medium (Cat. 18061) in SepMate™ tubes (Cat. 85450) per manufacturer protocol (StemCell Technologies Canada Inc.). CD4+ T cells were isolated by immunomagnetic negative selection using the EasySep™ Human Naïve CD4+ T Cell Isolation Kit II (Cat. 17555, StemCell Technologies Canada Inc.). Antibodies and soluble cytokines were acquired from Biolegend. Purified CD4+ naïve T cells were seeded at 225,000 cells/mL in anti-CD3ε-coated flasks (Cat. 317326) with soluble anti-CD28 (Cat. 302902), soluble anti-IFN-γ (Cat. 506531) and recombinant growth factors IL-7 (Cat. 581902), IL-4 (Cat. 574002), and TGF-β (Cat. 781802) in two phases, adapted from Pham’s protocol (20) and Bi *et al.*’s findings regarding the influence of IL-7 on Th9 differentiation (17). One triplicate of wells had CD4+ cells cultured only with anti-CD3ε and anti-CD28 as a “Th0” control. Media for Th9 derivation culture consisted of Roswell Park Memorial Institute (RPMI) 1640 medium (Cat. 11875093, ThermoFisher Scientific) with 10% FBS (Cat. CA76327-086, VWR), 1% penicillin/streptomycin (Cat. 15140122, ThermoFisher Scientific), 1% sodium pyruvate (Cat. 11360070, ThermoFisher Scientific), 1% MEM-non essential amino acids (Cat. 11140076, ThermoFisher Scientific), 0.5% HEPES (Cat. 15630080, ThermoFisher Scientific), 0.5% L-Glutamine (Cat. 25030081, ThermoFisher Scientific), and 0.05% β-mercaptoethanol (Cat. 31350010, ThermoFisher Scientific). After 5 days in culture, cells were stimulated with phorbol myristate acetate (PMA) (Cat. 74042, StemCell Technologies Canada, Inc.) and ionomycin (Cat. 73722, StemCell Technologies Canada, Inc.) for 5 hours before collecting cells and supernatant. PBS was added as control to the unstimulated cells.

### Syngeneic mouse model of endometriosis

*In vivo* experiments were done using our established syngeneic mouse model of endometriosis (48, 49). C57BL/6 mice (n= 10) were acquired from Charles River Laboratories (strain code 027) and housed in cages of four to five mice. Experiments were approved by the Queen’s Institutional Animal Care Committee. Donor mice were euthanized with inhalation of 5% isofluorane for three minutes followed by cervical dislocation. Uterine horns were removed and placed in PBS. Using a dermal biopsy punch, 3mm^3^ fragments of endometrial tissue were taken and kept on ice in PBS until surgically implanted in recipient mice. Prior to surgery, recipient mice were anesthetized in a vaporizer with 3.5% isofluorane. A small incision was made in the abdomen of each recipient mouse and two 3mm^3^ fragments of donor mouse endometrium were grafted in the left side of the peritoneal cavity of recipient mice using VetBond™ adhesive (Cat. 1469SB, 3M), while sham surgeries involved abdominal incision and suture without implantation of fragments. Postoperative fluid therapy and analgesics were given for 3 days following surgery and lesions were allowed to establish for 14 days. Peritoneal lavage, spleen, and endometriotic lesions were harvested at endpoint.

### Evaluating Th9 presence in mouse model of endometriosis

To capture baseline presence of Th9 cells in a mouse model of endometriosis, mice were induced with endometriosis (n= 3) or received sham operation (n= 3). Endometriotic lesions were allowed to develop over 14 days, then animals were euthanized to harvest spleen and peritoneal immune cells by peritoneal lavage.

Flow cytometry was performed in two panels of markers to quantify Th9 cells and, separately, mast cells (MCs) and committed mast cell progenitor (MCcp) presence in mouse splenocyte and peritoneal immune cell populations of mice induced with endometriosis compared to sham-operated mice. Spleens were mechanically digested through a 70 µm strainer into a single cell suspension in RPMI culture medium (Cat. 11835030, ThermoFisher Scientific) with 10% FBS. Peritoneal cells and splenocytes were counted by trypan blue exclusion in Countess 3 FL automated cell counter (ThermoFisher Scientific). To detect Th9 cells, peritoneal and spleen cells were stained with fluorescent antibody markers CD45—Pacific Blue, CD4— APC, IL-4Rα—PE/Cy7, and IRF4—PerCP/Cy5.5, as well as CD8α—FITC, CD14—FITC, CD19—FITC and Zombie Aqua fixable viability dye (Biolegend, San Diego, CA). Cells that were CD45^+^CD4^+^IL-4Ra^+^IRF4^+^CD8a^-^CD14^-^CD19^-^ were considered Th9 cells. In the second panel, to detect MC and MCcp, peritoneal cells were stained with fluorescent antibody markers CD45—Pacific Blue, FCERIα—PE, CD117—Brilliant Violet 650, CD11b—FITC, integrin β-7—APC, IL-9R—PE/Cy7 and Zombie Aqua fixable viability dye (Biolegend, San Diego, CA). MCs are identified as CD45^+^FCERIα^+^CD117^+^CD11b^-^, while MCcp are identified as CD45^+^CD117^+^integrin β-7^+^SSC^lo^.

### Differentiation of mouse Th9 cells in culture

Spleen and lymph nodes of C57/Bl6 mice (n= 3) (Charles River Laboratories) were harvested and mechanically digested through a 70µm strainer into a single cell suspension. Naïve CD4+ T cells were separated by negative selection using the EasySep™ Mouse Naïve CD4+ T Cell Isolation Kit (Cat. 19765, StemCell Technologies Canada Inc.) according to manufacturer protocol. Soluble cytokines and antibodies were obtained from Biolegend per their Th9 Polarization Activation Bundle (Th9 Polarization of Mouse CD4+ Cells Protocol, Biolegend).

Naïve CD4+ T cells were cultured in flasks pre-treated with plate-bound anti-CD3ε (Cat. 100340). Cultures were treated with a differentiation cocktail consisting of 5 μg/mL anti-CD28 (Cat. 102116), 20 ng/mL IL-4 (Cat. 574302), 2 ng/mL human TGF-β (Cat. 781802) and 10 μg/mL anti-IFN-γ (Cat. 505834) for 72 hours, followed by 10 ng/mL human IL-2 (Cat. 575402), 20 ng/mL mouse IL-4, and 1 ng/mL human TGF-β for 48 hours in 3X original media volume as per Pham’s published protocol (20). One triplicate was treated with 1x10^-6^ M E2 (Cat. E2758, Sigma Aldrich) to gauge impact of estrogen on Th9 differentiation. At the end of the 48-hour phase, CD4^+^ T cells were then activated *in vitro* with 50 ng/mL PMA and 750 ng/mL ionomycin for 6 hours before analysis by flow cytometry. One triplicate was treated with 1 µM monensin-containing protein transport inhibitor GolgiStop (Cat. 554724, BD Biosciences) to evaluate its impact on phenotype representation after PMA/ionomycin stimulation. Cells were stained with the following antibodies obtained from Biolegend: Pacific Blue—CD45, FITC—CD8α, CD14, CD19, APC—CD4, PerCP/Cy5.5—IRF4, PE-Cy7—IL-4Rα, Zombie Aqua fixable viability dye. Cells that were CD8α ^—^CD14^—^CD19^—^CD4^+^IRF4^+^IL-4Rα^+^ were considered Th9 cells.

### Purification and adoptive transfer of GFP+ Th9-like lymphocytes in a mouse model of endometriosis

Spleen and lymph nodes of donor GFP+ C57/Bl6 mice (C57BL/6-Tg(UBC-GFP)30Scha/J, strain # 004353, Jackson Laboratory) (n= 5) were dissected, mechanically digested and passed through a 70µm strainer to obtain a single cell suspension. CD4+ cells were isolated and cultured as described above. Before adoptive transfer, Th9 cells were purified by two positive selection immunomagnetic separation kits. StemCell EasySep™ Release Mouse Biotin Positive Selection Kit (Cat. 17655) and PE Positive Selection Kit (Cat. 17656) were used per manufacturer protocols to positively select IL-4Rα+ and subsequently TGF-β1 RII+ cells. As IL-4Rα and TGF-β1 RII are not concurrently expressed in any other T cell subsets other than Th9 cells, this allowed us to obtain a purified Th9 population. A PE-conjugated TGF-β1 RII antibody (Cat. FAB532P, R&D Systems) was used with the PE positive selection kit. A purified IL-4Rα antibody (Cat. 144801, Biolegend) was biotinylated using Abcam’s biotinylation kit (Cat. AB201795) per manufacturer protocol and was used to select IL-4Rα+ cells in the biotin positive selection kit. On day 0, endometriosis was induced in recipient C57/Bl6 mice (n= 10) with two 3mm^3^ endometrial fragments as detailed previously. On day 7, endometriosis-induced mice (n= 5) each received a single intraperitoneal injection of approximately 274,000 GFP+, *in vitro-*expanded, purified Th9-like lymphocytes suspended in 100µL of PBS. Concurrently, a group of endometriosis-induced mice (n= 5) received intraperitoneal injections of 100µL PBS as a control. Blood was collected at day 7 and day 14 to measure plasma cytokine levels before and after adoptive transfer (Figure 5A). Specifically, using submandibular vein puncture, 100 µl of blood was collected in EDTA-coated collection tubes and blood was allowed to coagulate on ice for 2 hours. Blood samples were centrifuged at 3000g for 15 minutes at 4 °C and separated plasma was aspirated before being diluted 1:2 in preparation for multiplex cytokine analysis.

Plasma cytokines, chemokines and growth factors were analyzed using a commercially available 32-plex panel (MD-32, Eve Technologies, Calgary, AB, Canada). Animals were euthanized on day 14 as previously described in order to collect peritoneal fluid (lavage), lesions, spleen, and uterus. Specifically, one lesion was preserved in freshly prepared 4% paraformaldehyde for 24-hour fixation at 4 °C while the other lesion was snap-frozen in liquid nitrogen and stored in -80 °C to be processed for RNA extraction.

### Evaluation of differentially expressed genes using NanoString transcriptomic analysis in endometriotic lesions from mice with or without Th9 adoptive transfer

Mouse endometriotic lesions were homogenized using ceramic beads (Cat. #13113-50, Qiagen N.V., Hilden, Germany) in the Omni Bead Ruptor 24 (model # 19-010, Omni International Inc). Total RNA was extracted and purified from lesion tissue lysate using Norgen Total RNA + Micro Isolation Kit (Cat. # 48500) as per manufacturer instructions. RNA concentrations were normalized to 120 ng/µl, with RNA concentration and purity verified using NanoDrop 2000 spectrometer (Thermo Fisher Scientific). Using the Mouse Fibrosis V2 Panel (NanoString, Seattle, WA, USA), 760 genes were evaluated in mouse lesion RNA. Hybridization, sample processing and data collection was conducted by the Ontario Institute of Cancer Research (Toronto, ON, Canada). Briefly, samples were hybridized with probes over an 18-hour incubation, then loaded into the nCounter cartridge, and digital counts were obtained across 280 fields of view (FOV) using the nCounter Digital Analyzer. Data were normalized to internal controls and housekeeping genes to ensure accurate quantification across samples using nSolver software. Any housekeeping genes with an average count <100 were removed and any genes with a maximum count <20 were not included in analysis. Evaluation of differentially expressed genes was conducted using nSolver software (version 4.0), in which heat maps were generated to display data in unsupervised clusters. Advanced analysis was conducted to detect patterns of differential expression in genes associated with specific functions, including Th subset differentiation and type 2 inflammatory responses. Through nSolver’s advanced analysis functions, Pathview analysis was conducted to illustrate significant changes to activity of specific pathways related to focal adhesion, chemokine signalling pathways, cytokine-cytokine receptor interaction, and PI3K-AKT signalling.

### Immunohistochemistry of mouse endometriotic lesions

Mouse endometriotic lesion tissues were fixed for 24 hours in 4% paraformaldehyde at 4 °C before being transferred to 70% ethanol. Tissues were then dehydrated and paraffinized over 11 hours and embedded in paraffin blocks. Sections were cut at 5µm and mounted on glass slides for immunohistochemistry staining using a Leica Bond RX automated stainer (Leica Microsystems). Sections underwent citrate-based epitope retrieval for 20 minutes before 15-minute incubation with primary antibodies for proliferation marker Ki67 (1:3000, Cat. ab15580, Abcam) or angiogenic marker CD31 (1:300, Cat. 77699S, New England Biolabs). Leica BOND Polymer Refine Detection kit (Leica Microsystems) was used for 3,3’-Diaminobenzidine (DAB) chromogen detection and hematoxylin counterstain. Slides were scanned at 40X magnification in an Olympus VS120 high resolution slide scanner (Olympus Life Science). Images were analyzed with algorithms designed in HALO AI (Indica Labs, Albuquerque, New Mexico, USA). Using the HALO tissue classifier feature, tissue was classified into slide glass (empty area), stroma, epithelium, vasculature, and glandular lumen. An area quantification algorithm was designed to analyze CD31 staining of vascular tissue areas, while a cytonuclear algorithm was designed to analyze Ki67+ cells. Percent positive cells or positive area from each tissue type were counted by their respective algorithms and data was imported to GraphPad Prism for statistical analysis.

### Immunofluorescence of mouse endometriotic lesions

Mouse endometriotic lesion tissues were sectioned at 5µm and mounted on glass slides before undergoing deparaffinization and citrate-based heat antigen retrieval for 20 minutes. Slides were permeabilized in PBS-T for 10 minutes followed by peroxidase suppression for 10 minutes (Cat. 35000, ThermoFisher Scientific). After 1 hour blocking in 5% bovine serum albumin PBS-T solution, slides were incubated with primary antibodies rabbit anti-mouse IL-9 (1:200, Cat. Ab203386, Abcam), rat anti-mouse CD4 (1:50, Cat. MA1146, ThermoFisher Scientific), and chicken anti-GFP (1:100, Cat. A10262, ThermoFisher Scientific) overnight at 4 °C. After three 10-minute washes with PBS-T, slides were incubated with secondary antibodies donkey anti-rabbit Alexa Fluor 568 (1:400, Cat. A10042, ThermoFisher Scientific), donkey anti-rat Alexa Fluor 647 (1:400, Cat. A78947, ThermoFisher Scientific), and goat anti-chicken Alexa Fluor 488 (1:200, Cat. A11039, ThermoFisher Scientific) for two hours in the dark at room temperature. Finally, slides were washed thrice with PBS-T at 10 minutes per wash, then twice with PBS at 5 minutes per wash, before mounting with DAPI-containing SlowFade Glass mounting medium (Cat. S36920, Thermo Fisher Scientific). Slides were scanned using a Leica Mica confocal microscope and visualized using the LAS X Life Science Microscope Software (Leica Microsystems).

## Statistical analyses

For cytokine analyses, flow cytometric population comparison, and tissue microarray IL-9 stain comparison, one-way ANOVA tests were conducted in GraphPad Prism, version 10. For any comparison between only two groups (endometriosis vs. sham, adoptive transfer vs. PBS control) a student’s t-test was conducted in GraphPad Prism. Significance was considered p≤0.05. Outlier tests were conducted on each set of data (ROUT Q= 1%) and all statistical tests were performed on cleaned data.

## Data availability

Data sets are available from authors upon request.

## Author contributions

AM and CT designed research studies. AM, KBZ, DJS, PY, HL, DVH, and AKR conducted experiments. AM acquired data, analyzed data, and wrote the manuscript. AM, KBZ, DJS, HL, and CT revised the manuscript. CT provided reagents and equipment.

## Acknowledgements

This work was supported by grants from Canadian Institute of Health Research and Natural Sciences and Engineering Research Council of Canada.

## Supplemental figure

**Supplemental figure 1.**
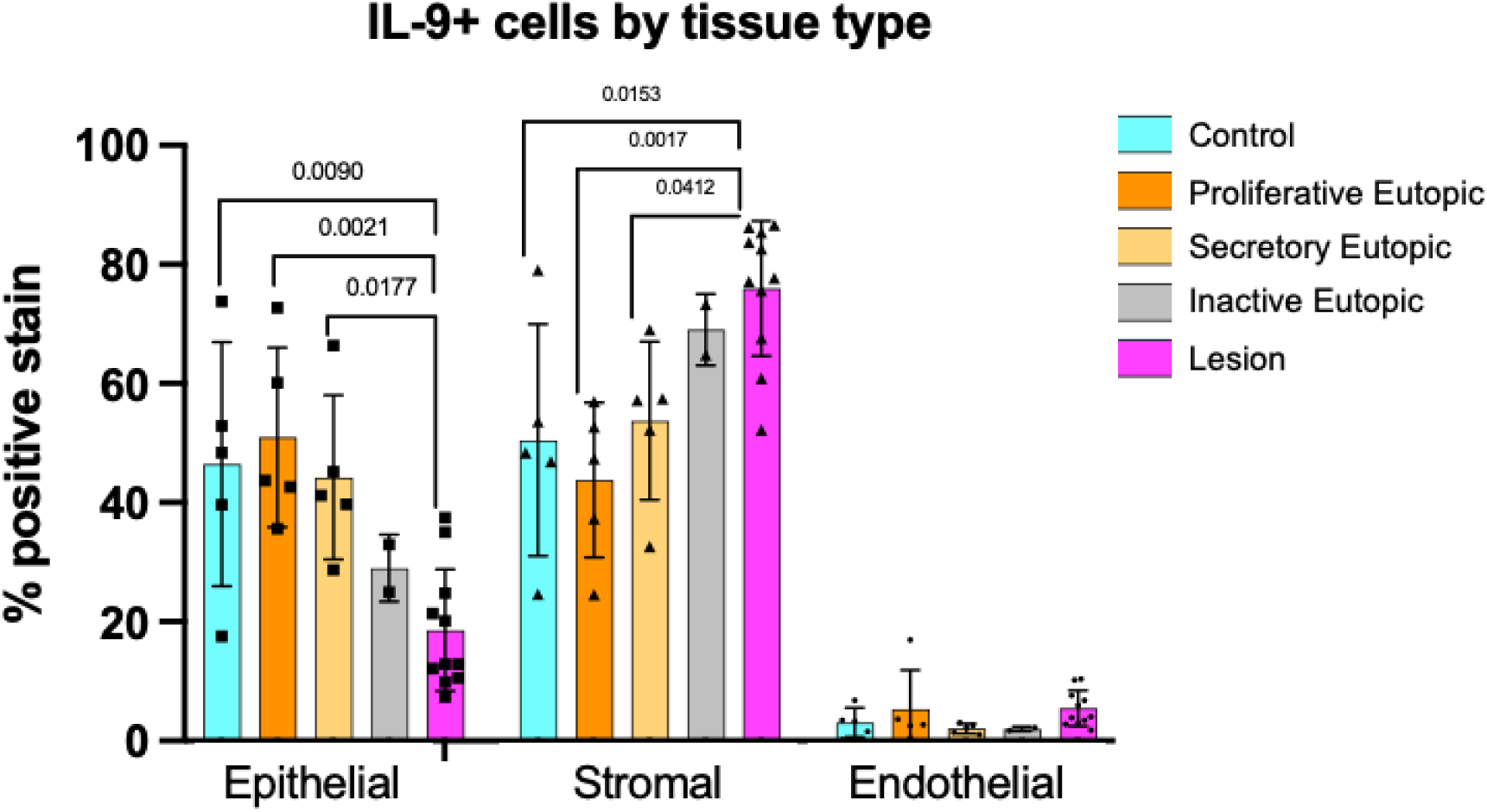
Stromal proportion of IL-9+ cells is significantly higher in lesion samples, epithelial proportion significantly lower. Out of IL-9+ cells, the portion represented by stromal cells was significantly higher for the lesion sample group (n= 11) as compared to control endometrium (n= 5), proliferative eutopic endometrium (n= 5), and secretory eutopic endometrium (n= 5). Inactive endometrium appears to be most similar to lesion tissue in terms of IL-9+ distribution.

